# Varying co-factor requirements for MAML1-dependent transcription at different Notch-responsive target genes

**DOI:** 10.1101/2020.01.13.905612

**Authors:** Julia M. Rogers, Bingqian Guo, Emily D. Egan, Jon C. Aster, Karen Adelman, Stephen C. Blacklow

## Abstract

Mastermind proteins are required for transcription of Notch target genes, yet the molecular basis for mastermind function remains incompletely understood. Previous work has shown that Notch can induce transcriptional responses by binding to promoters, but more often by binding to enhancers, with *HES4* and *DTX1* as representative mammalian examples of promoter and enhancer responsiveness, respectively. Here, we show that mastermind dependence of the Notch response at these loci is differentially encoded in Jurkat T-ALL cells. Knockout of Mastermind-like1 (MAML1) eliminates Notch responsive activation of both these genes, and reduced target gene expression is accompanied by a decrease in H3K27 acetylation, consistent with the importance of MAML1 for p300 activity. Add-back of defined MAML1 constructs in knockout cells identifies residues 151-350 of MAML1 as essential for expression of either Notch-responsive gene. Fusion of the Notch-binding region of MAML1 to the HAT domain of p300 rescues expression of *HES4* but not *DTX1*, suggesting that an additional activity of MAML1 is needed for gene induction at a distance. Together, these studies establish the functional importance of the MAML1_151-350_ region for Notch-dependent transcriptional induction, and reveal differential requirements for MAML1-dependent recruitment activities at different Notch responsive loci, highlighting the molecular complexity of NTC-stimulated transcription.

## Introduction

Notch proteins are receptors in a juxtracrine signaling pathway that influences numerous developmental patterning decisions^1,2^. Dysregulation of Notch signaling contributes to the pathogenesis of many human cancers, including T-cell acute lymphoblastic leukemia, a disease in which activating mutations of human Notch1 are well established oncogenic drivers^3–5^.

Notch signaling is initiated by ligand binding, which induces proteolytic liberation of the intracellular portion of Notch (NICD) into the cell, where it enters the nucleus to activate transcription. In the nucleus, it forms a Notch transcription complex (NTC) with the sequence specific DNA binding protein RBPJ (also known as CSL) and a transcriptional coactivator of the Mastermind-like (MAML) family^6–8^. The mechanisms by which NTC genomic binding activates transcription are an active area of study.

MAML proteins are required for transcription of mammalian Notch target genes, highlighting the importance of studying MAMLs to understand the transcriptional response to Notch activation^8^. In human MAML1, residues 13-74 directly interact with RBPJ and NICD, but the remainder of the 1016 amino acid long protein is poorly characterized^9^. C-terminal truncations of MAML1 have dominant-negative activity, presumably because these fragments bind the NTC but cannot interact with the other co-regulators that it is responsible for recruiting^8,10^. These findings have motivated further characterization of the function of the remaining C-terminal portion of MAML1, in order to identify what co-factors it recruits, and how these partners contribute to transcriptional activation.

Prior studies suggest that MAML1 has two regions that contribute to transcriptional activation. A GST fusion protein with the first region (TAD1), which includes residues 75-301, binds to p300 recovered from nuclear extracts, and a MAML1_1-301_ fragment, which includes both the region required for NTC entry and TAD1, is sufficient to stimulate transcription from an *in vitro* chromatinized template^11^. However, it is not clear that MAML1 is directly responsible for recruiting p300 to the genome, as ChIP-PCR of the *HES1* locus showed p300 at the promoter in the absence of Notch activation and before NTC binding^12^. In addition, a similar fragment, MAML1_1-300,_ acts as a dominant-negative protein in reporter assays, further complicating interpretation of how p300 binding is related to transcriptional induction by NTCs^10^. The remaining C-terminal portion of MAML1 (TAD2) is not required for transcription *in vitro*, but is required in cells^11^. One line of investigation has suggested that this part of MAML1 is required for recruitment of cyclin C:CDK8 complexes (and thus, the kinase module of the mediator complex), thereby promoting phosphorylation of NICD, but how these activities are coordinated and how they contribute to target gene expression remain incompletely understood^12^.

Such mechanistic studies looking at regulation of Notch responsive genes have centered on canonical targets such as *HES1*, which is regulated by promoter proximal Notch binding^12^. Most cell type-specific Notch-responsive genes, however, are regulated by binding of Notch transcription complexes to distal enhancers^13–16^. The conflicting requirements reported for different MAML1 activities *in vitro*, in cells, and in organisms highlight the importance of studying Notch function in physiologically relevant cellular systems, and suggest that more expansive approaches could uncover different roles for MAML binding at other genomic loci.

Here, we have interrogated the function of MAML1 fragments in regulating Notch-dependent gene expression in Jurkat T-ALL cells, examining established target genes regulated by either promoter proximal Notch binding or distal enhancer binding. We greatly extend prior work by creating MAML1-knockout cells and by using them to show that a form of MAML1 lacking residues 151-350, a region that overlaps TAD1, is defective in rescuing Notch target gene expression in MAML1 knockout cells. Furthermore, we establish that reduced target gene expression is accompanied by a decrease in H3K27 acetylation, consistent with the importance of the excised 151-350 region for p300 recruitment. However, when the Notch binding region of MAML1 is fused to the HAT domain of p300, the fusion protein is able to rescue expression of *HES4*, the proximal-promoter regulated gene, but not *DTX1*, the enhancer regulated Notch target gene, suggesting that an additional activity of MAML1 is needed for gene induction at this locus. Together, these studies establish the functional importance of the MAML1_151-350_ region for Notch-dependent transcriptional induction, and reveal differential requirements for MAML1-dependent recruitment activities at different Notch responsive loci, highlighting the molecular complexity of NTC-stimulated transcription.

## Results

### Jurkat cells exhibit dynamic Notch responsiveness dependent on MAML1

To investigate MAML structure-function relationships, we turned to the Jurkat T-ALL cell line because it has a juxtamembrane insertion in Notch1 that renders the protein constitutively active, but does not depend on Notch activity for survival^5,17^. Furthermore, the constitutive activity of the mutated Notch1 protein in these cells relies on gamma-secretase cleavage for release from the membrane, allowing us to use a well-established gamma secretase inhibitor (GSI) washout assay^18^ to toggle between Notch-off (GSI) and Notch-on (washout) states. With this approach, we confirmed that *HES4* and *DTX1* are direct Notch target genes in these cells by RT-qPCR (Figure 1a). Expression of each of these genes is suppressed by GSI treatment and restored upon inhibitor washout, a characteristic feature of Notch target genes.

**Figure 1:**
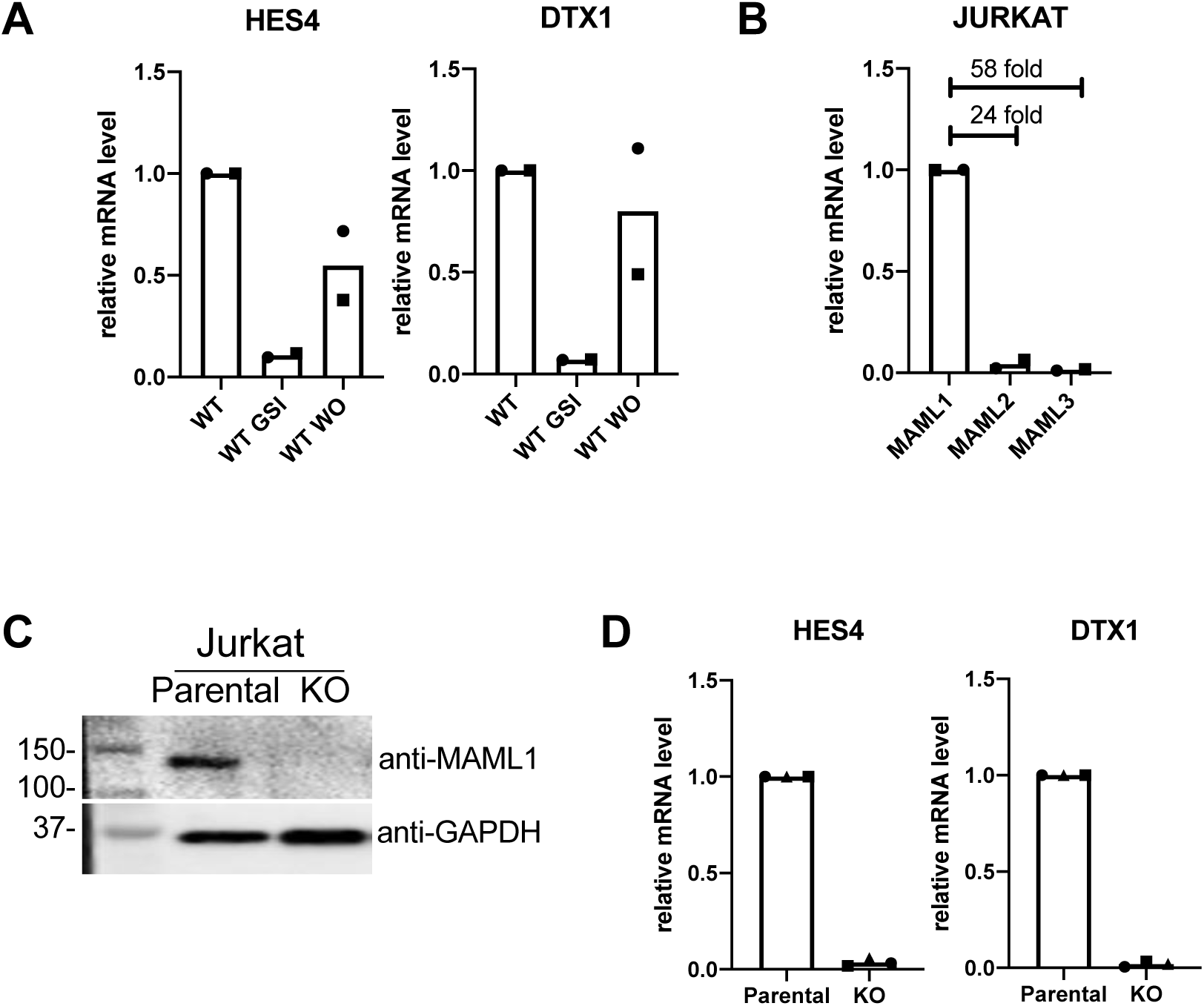
Jurkat cells are dependent on MAML1 for Notch stimulated transcription. (a) *HES4* and *DTX1* are Notch-responsive genes in Jurkat cells. RT-qPCR results in Jurkat cells growing in RPMI media, compared to media with gamma secretase inhibitor (GSI), and media after GSI washout (WO). Bars indicate the mean of two biological replicates. Data points show the average of technical replicates from each biological replicate, with each of the two biological replicates indicated by a different point shape. (b) Comparison of MAML1, MAML2, and MAML3 transcript abundance in Jurkat parental cells, as determined by RT-qPCR. Bars indicate the mean of two biological replicates. Data points show the average of technical replicates from each biological replicate, with each of the two biological replicates indicated by a different point shape. (c) Western blot for MAML1 protein in parental and MAML1 KO Jurkat cells. (d) Expression of Notch target genes *HES4* and *DTX1* in parental and MAML1 KO Jurkat cells, as determined by RT-qPCR. Bars indicate the mean of three biological replicates. Data points show the average of technical replicates from each biological replicate, with the three biological replicates indicated by a different point shape.

Next, we determined the transcript abundance of the three human MAML proteins in Jurkat cells by RT-qPCR. MAML1 transcripts are much more abundant in these cells than are MAML2 and MAML3 (Figure 1b), consistent with prior studies implicating MAML1 as the most important member of this family in lymphocyte development *in vivo*^19^. Therefore, we created a clonal MAML1 knock-out (KO) cell line using CRISPR-Cas9 genome editing (Figure 1c), to enable subsequent add-back studies for structure-function analysis. Consistent with the conclusion that *HES4* and *DTX1* are indeed Notch target genes in this cell context, their expression is greatly suppressed in the KO line (Figure 1d,e).

### Amino acids 151-350 of MAML1 are essential for Notch activation

In order to elaborate structure-function relationships, we created a series of truncation and internal deletion mutations of MAML1. We first assessed the conservation, predicted disorder, and charge distribution in MAML1. Residues 13-74 of MAML1 interact directly with Notch1 and RBPJ, forming the Notch transcriptional activation complex^9,20,21^. This N-terminal region is also more highly conserved and predicted to be more ordered than the remainder of the protein (Figure 2). There are two other acidic segments that are both conserved and predicted to be ordered, 263-276 and 990-1016. The C-terminal segment has been reported to associate with the human papilloma virus E6 protein, but precise functions for these regions have otherwise not been assigned^22^. The remainder of the MAML1 sequence is predicted to be disordered (Figure 2, middle panel).

**Figure 2:**
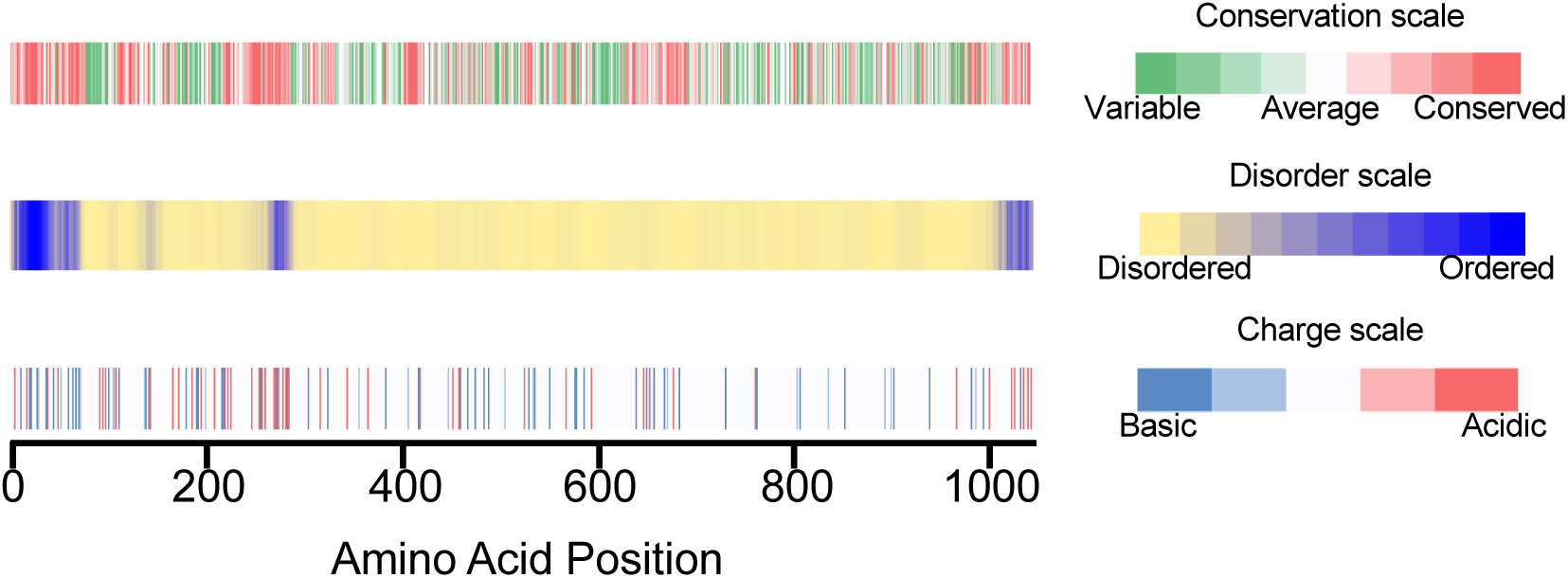
MAML1 sequence features inform regions to assay. Top: Conservation of MAML1 sequence, measured by Conseq. Middle: Predicted disorder in the MAML1 sequence, predicted using Disopred3. Bottom: Amino acid charge throughout the MAML1 sequence.

We created several constructs to assess the function of regions of MAML1 (Figure 3a). All constructs retained the N-terminal 150 amino acids in order to span the nuclear localization signal (NLS) and the Notch interacting region, and included a C-terminal GFP tag and a mutated PAM sequence to avoid recognition by Cas9 in the add-back cell lines.

**Figure 3:**
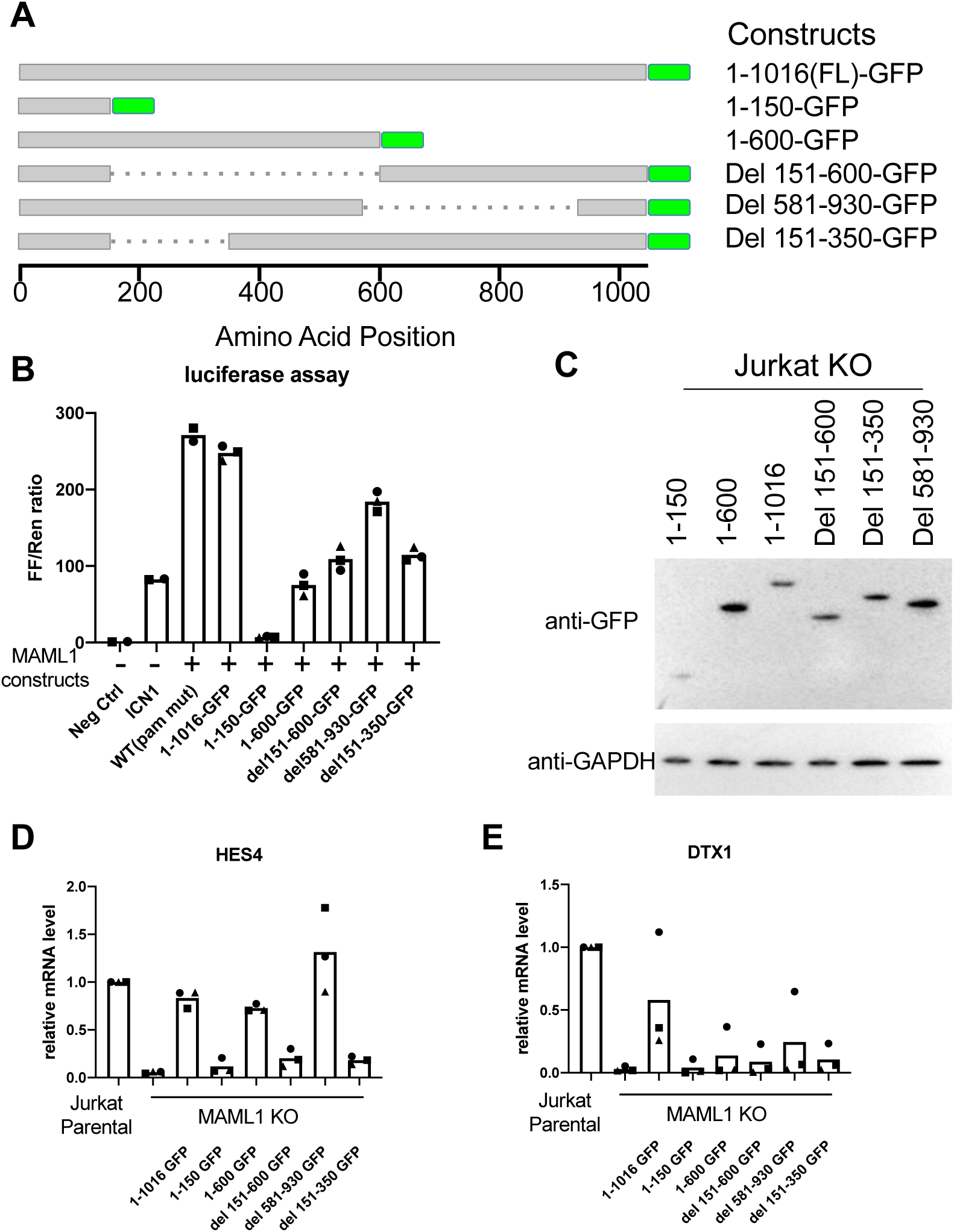
Region 151-350 of MAML1 is required for Notch target gene expression. (a) Constructs assayed in add-back experiments. All constructs contain the MAML1 NLS, and a C-terminal GFP tag. (b) Luciferase assay performed in U2OS cells shows the function of different MAML1 constructs in enhancing Notch-mediated transcription from a Notch reporter. Bars indicate the mean of biological replicates. Data points show the average of technical replicates from each biological replicate, with each biological replicate indicated by a different point shape. (c) Stable cell lines expressing various MAML1 truncations. Expression of the MAML1 constructs is detected by Western blot with an anti-GFP antibody. (d-e) RT-qPCR of Jurkat parental cells, KO, and add-back lines with the various MAML1 constructs is shown for (d) *HES4* and (e) *DTX1*. Bars indicate the mean of 3 biological replicates. Data points show the average of technical replicates from each biological replicate, with each of the three biological replicates indicated by a different point shape.

These constructs were first tested in a Notch luciferase reporter assay. The C-terminal GFP tag did not reduce activity compared to untagged protein (Figure 3b). MAML1_1-150_ acts as a dominant negative construct, as has been reported for other N-terminal MAML1 fragments, which bind Notch and RBPJ, but are unable to stimulate transcription^10^. The other constructs all reduced activity to varying levels: MAML1_1-600_ by 70%, MAML1_Δ151-600_ and MAML1_Δ151-350_ by over 50%, and MAML1_Δ581-930_ by 25%. The dramatic difference in the behavior of MAML1_1-600_ and MAML1_Δ581-930_ suggests an important role of the C-terminal acidic region, which is present in the more active MAML1_Δ581-930_, but absent in MAML1_1-600_. The fact that MAML1_Δ151-600_ and MAML1_Δ151-350_ behave similarly indicates that the removal of residues 151-350 is primarily responsible for the reduced activity.

We then created stable cell lines expressing these constructs (Figure 3c, Figure S1a, Figure S1b) in the MAML1 KO background, and measured expression of Notch target genes (Figure 3d-e). All constructs that lack positions 151-350 are unable to rescue either *HES4* or *DTX1* expression. MAML1_1-600_ rescues expression to varying amounts (∼14% for *DTX1* to 72% for *HES4*). MAML1_Δ581-930_ rescues expression for *HES4*, but not for *DTX1*, suggesting different requirements for the C-terminal portion of MAML1 for different genes.

### Defective activation by MAML1_Δ151-350_ is not due to a change in activated Notch1 stability

As all segments of MAML1 missing amino acids 151-350 impacted function, we focused on how this segment contributes to gene expression. We assessed the level of active, intracellular Notch in parental, KO, full length add-back, and MAML1_Δ151-350_ (“del”) add-back cell lines (Figure 4, Figure S2). In the parental line, NICD1 undergoes substantial phosphorylation, as judged by the ability of lambda phosphatase to collapse the protein band on Western blot to a more focused, faster migrating species. The NICD1 in the MAML1 KO cells migrates similarly to the lambda-phosphatase treated protein, suggesting that MAML1 is required for most NICD1 phosphorylation in these cells. Phosphorylation of NICD1 is rescued by both the full-length and del proteins, although the del protein appears to show differences in the proportions of various NICD1 phosphorylated species. Examination of the NICD1 abundance after phosphatase treatment shows, however, that the overall abundance of Notch protein is not lower in the “del” line compared to WT, indicating that this form of MAML1 is not deficient in transcription because of effects on NICD1 overall abundance or stability.

**Figure 4:**
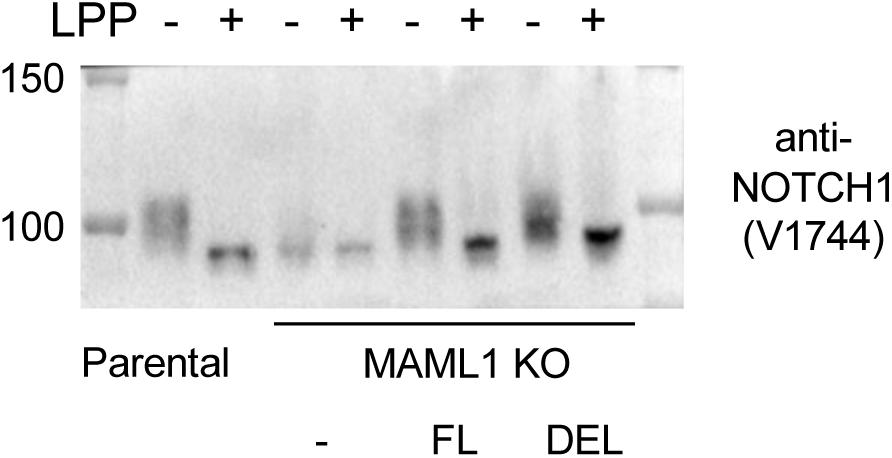
MAML1 rescue constructs restore normal NICD1 abundance. Western blot for activated Notch (NICD1) in parental Jurkat, MAML1 KO, MAML1_1-1016_ add-back and MAML1_Δ151-350_ add-back cell lines. Lanes marked “+” were treated with lambda phosphatase.

### MAML1_Δ151-350_ does not change the abundance of chromatin bound Notch1, but does reduce H3K27 acetylation around Notch target genes

We then assessed chromatin-bound Notch protein amounts at the *HES4* and *DTX1* Notch target genes in the four cell lines (parental, KO, full length and “del” add-back) by ChIP-qPCR (Figure 5a,b). As anticipated based on the reduced amount of total Notch1 protein in the MAML1 KO cells, MAML1 KO reduced Notch abundance at both the *HES4* and *DTX1* binding sites. Both full-length and “del” add-back rescued Notch genomic binding levels, indicating that residues 151-350 of MAML1 are not responsible for stabilizing Notch bound to its response elements on DNA. We next measured H3K27ac levels surrounding the Notch binding sites (Figure 5a,c,d). MAML1 KO cells have reduced H3K27ac at Notch targets, which is rescued by full-length MAML1, but not by MAML1_Δ151-350_, indicating that this segment promotes histone acetylation surrounding Notch genomic responsive elements.

**Figure 5:**
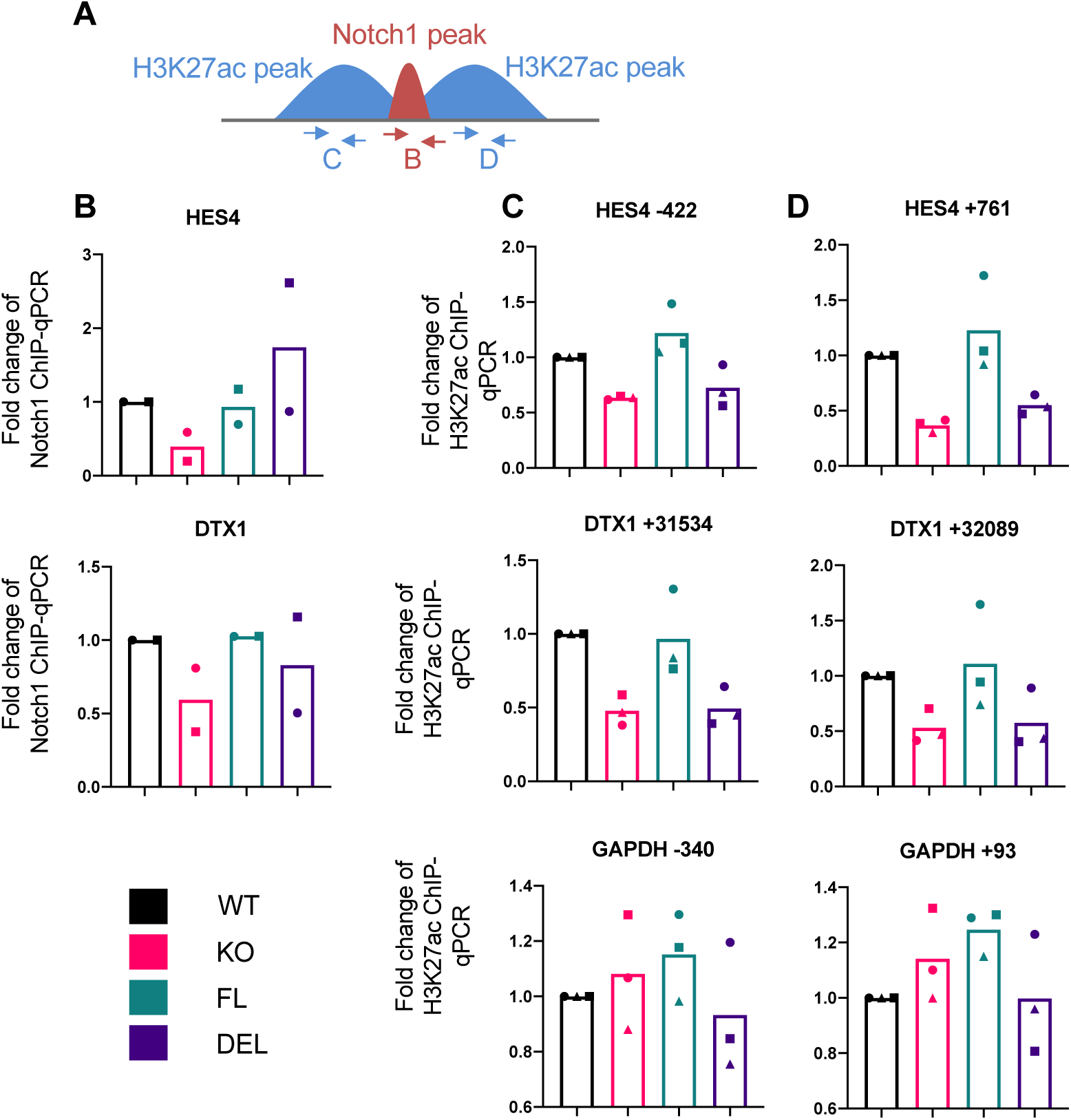
MAML1_Δ151-350_ cannot rescue H3K27ac around Notch binding sites. (a) Scheme showing primer placement for Notch1 and H3K27ac ChIP-qPCR experiments. (b) Notch1 binding at regulatory regions for *HES4* and *DTX1*, as measured by Notch1 ChIP-qPCR. Bars indicate the mean of two biological replicates. Data points show the average of technical replicates from each biological replicate, with each of the two biological replicates indicated by a different point shape. (c,d) H3K27ac signal flanking the Notch1 binding sites is shown, as measured by ChIP-qPCR. H3K27ac flanking the GAPDH promoter is shown as a non-Notch responsive control. Bars indicate the mean of three biological replicates. Data points show the average of technical replicates from each biological replicate, with each of the three biological replicates indicated by a different point shape.

### MAML1 1-150 p300HAT fusion partially rescues target gene expression

To determine if gene expression could be restored by artificial recruitment of histone acetyltransferase activity, we created a fusion of MAML1 to the p300 histone acetyltransferase (HAT) domain. We linked the N-terminal 1-150 amino acids of MAML1, containing the NLS and Notch binding region, to the p300 HAT domain (Figure 6a). We created a stable cell line in which this fusion was expressed in MAML1 KO cells, and measured Notch target gene expression. This fusion rescued 77% of *HES4* expression. Importantly, this rescue of target gene expression was suppressed in the presence of GSI, indicating that rescue by the MAML1-HAT fusion is dependent on the presence of intracellular Notch (Figure 6b). The MAML1-HAT fusion, however, did not measurably rescue expression of DTX1 (Figure 6c).

**Figure 6:**
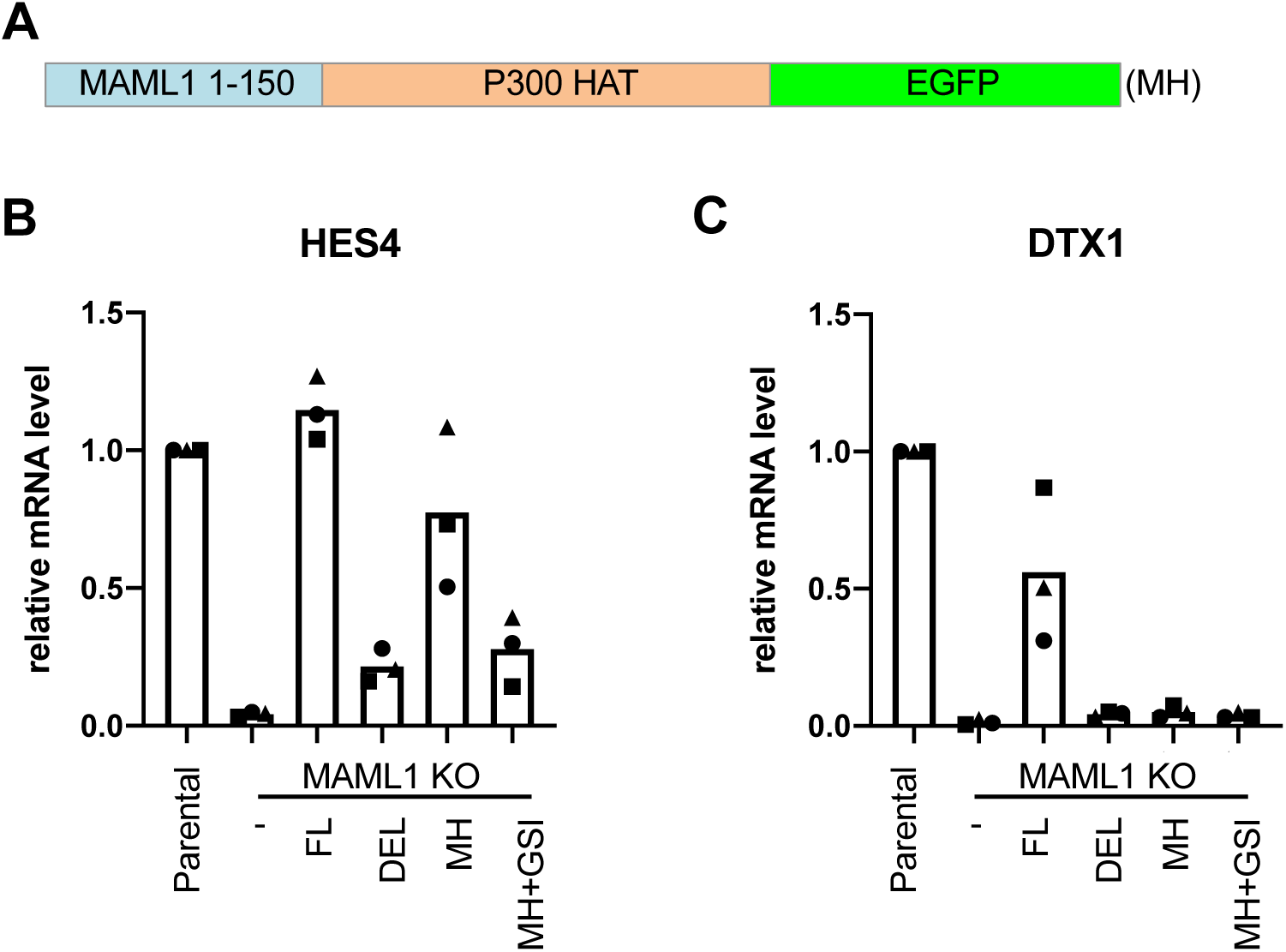
A MAML1-p300 fusion protein can rescue expression of some Notch targets. (a) Scheme of the designed MAML1-HAT (MH) fusion protein. (b-d) RT-qPCR of Notch responsive genes *HES4* (b) and *DTX1* (c) in MAML1 KO Jurkat cells expressing FL, MAML1_Δ151-350_, MH, or MH in the presence of a gamma secretase inhibitor (GSI). Bars indicate the mean of 3 biological replicates. Data points show the average of technical replicates from each biological replicate, with the three biological replicates indicated by a different point shape.

## Discussion

In this work, we have shown the requirement of amino acids 151-350 for MAML1 function in both reporter assays and at endogenous loci in Jurkat cells. This finding is consistent with previous work identifying TAD (residues 75-301) as required for transcription. Interestingly, previous work has shown that the p300 interaction region of MAML1 residues in a proline rich motif within amino acids 81-87, which is retained in our MAML1_Δ151-350_ construct. Because MAML1_Δ151-350_ cannot support histone acetylation at Notch responsive binding sites, our data suggest that binding of MAML1 to p300 through the previously defined site (*i.e.* residues 81-87) is not sufficient for it to stimulate histone acetylation^23^, and that additional segments of MAML1 beyond the previously reported p300 binding site are required for recruiting p300 acetyltransferase activity. Interestingly, lysine residues in the 151-350 region of MAML1 can be acetylated by p300, suggesting the existence of a more complex interplay between MAML1-p300 binding and enzymatic activity than previously appreciated^23^.

We also showed that direct fusion of the MAML1 N-terminus to the p300 HAT domain could rescue expression of *HES4*, but not *DTX1. HES4* is regulated by promoter-proximal NTC binding, while *DTX1* is regulated by an intronic enhancer approximately 31 kB downstream. Previous studies tethering the p300 HAT domain to dCas9 have also found that gene activation through recruitment of p300 to enhancers is less robust than activation from promoters^24^. It is intriguing to consider how different Notch-responsive genes have encoded different requirements for transcriptional co-factors. *DTX1* is more dependent on the C-terminal regions of MAML1 than is *HES4*, highlighting that while promoter proximally regulated genes may be primarily dependent on p300 recruitment and activation, distally regulated genes may depend on recruitment of other co-activators by the C-terminal end of MAML1. Future work will be required to identify additional MAML1-dependent co-factors, define how they interact with MAML1 and establish how they contribute to expression of Notch target genes.

## Methods

### Cell culture

Jurkat cells were maintained in RPMI with L-glutamine media (Corning) + 10% FBS (Gemini Bio-Sciences) and Penicillin-Streptomycin (Gibco). U2OS and 293T cells were maintained in DMEM (Corning) + 10% FBS (Gemini Bio-Sciences) and Penicillin-Streptomycin (Gibco).

### GSI Treatment and Washout

Gamma secretase inhibitor Compound E (Millipore Sigma) was dissolved in DMSO and diluted 1:1000 in RPMI media to a final concentration of 1 μM. Cells were treated with compound E or an equal quantity of DMSO as control for 8 or 16 h. Cells from compound E-treated and DMSO control-treated conditions were then recovered by centrifugation (300 *g*, 5min). Cells were then subjected to three cycles of resuspension in RPMI followed by recentrifugation to remove the GSI or DMSO. Cells were harvested for analysis 4 hours after GSI or DMSO washout.

### Cloning

A guide sequence (GCGGTCATGGAGCGCCTTCGC) for CRISPR targeting of MAML1 was cloned into the LentiV2 vector using the Esp3I restriction enzyme. The various MAML1 truncation constructs were subcloned into the MigR1 vector using restriction sites for XhoI and NcoI. The resulting cDNAs eliminated the IRES site for expression of GFP, and instead encoded MAML1-GFP fusion proteins directly.

The MAML1-HAT fusion protein was cloned into the MigR1 vector using the same sites, also generating a C-terminal GFP fusion protein.

### Cell lines

The MAML1 knockout cell line was generated by infecting Jurkat cells with the lentiviral CRISPR-Cas9 system to target MAML1. Infected cells expressing GFP underwent single cell sorting and clonal expansion before Western blot validation. MAML1 rescue cDNA constructs had their PAM sequences mutated to prevent Cas9 cleavage.

Viral particles were made by co-transfecting MigR1-MAML1 plasmid, VSV-G, and gag-pol vectors into 293T cells. Viruses were harvested after 48 h and filtered through a 0.45 μm membrane. Jurkat cells were then infected with virus by mixing 0.5 million cells with virus and 4 μg/mL polybrene. GFP-positive cells were sorted 3-4 days after infection using a FACS Aria sorter.

### Reporter assays

50000 U2OS cells were plated per well, in a 24-well cell culture plate. The next day, cells were transfected with 10 ng ICN1 in pcDNA, 50 ng MAML1 construct in MigR1 vector, 100 ng of reporter mix (98 ng TP1/2 ng PRLTK), and empty pcDNA vector to 250 ng, using PEI in a 1:3 DNA: PEI ratio. Three replicate transfections were performed per condition. Cells were harvested the next day, and luciferase assays were performed using the Dual-Luciferase Reporter Assay Kit (Promega), read out on a Turner Biosystems Modulus Microplate luminometer, with 2 or 3 technical replicates per transfection. Firefly to Renilla luminescence ratios were normalized to mean of empty vector control = 1.

### Western Blots

For Western blots, cells were pelleted, resuspended in 1X SDS running buffer at 10^6^ cells per 100 μL, boiled and run on a 4-20% gradient gel (Bio-Rad). MAML1 was detected with CST antibody D3K7B for Figures 1C and S1a, and CST antibody D3E9 for Figure S1b. GAPDH was detected with CST antibody 14C10, and GFP with CST antibody 2956.

750000 cells were harvested and resuspended in 40 μL lambda protein phosphatase reaction buffer (1X NEBuffer Pack for Protein MetalloPhosphatases (NEB), 1mM MnCl_2_ (NEB), 0.1% Igepal NP-40, protease inhibitor tablet). 1uL of lambda protein phosphatase (NEB) was added to samples, according to the Figure 4 legend. Samples were incubated at 30C for 30 min, 700 rpm, then 10 μL of 5X SDS loading buffer was added. Samples were run on 8% Tris-Glycine Novex gels (Invitrogen) at 150V for 2 hrs, until the 75 kDa marker was at the bottom of the gel. Samples were ponceau stained to verify equal loading, and blotted with CST V1744 activated Notch antibody (4147).

### RT-qPCR

RNA was harvested from 1,000,000 cells using the RNEasy Kit (Qiagen). 1 μg of total RNA was used as input for cDNA synthesis, using the iScript kit (Bio-Rad). qPCR was performed using PowerUp SYBR Green Master Mix (Thermo Fisher) in a 10 μL total reaction with 0.25 μM forward and reverse primers, with 2 technical qPCR replicates per condition. Primer sequences are listed in Supplementary Table 1. The standard thermocycler parameters suggested by the manufacturer were used, with a default melt curve step right the PCR run to confirm primer specificity. Expression was normalized to the average GAPDH expression for each condition. In order to show biological replicates on the same scale, within each replicate expression was normalized to WT Jurkat cell expression = 1.

### ChIP-qPCR

Chromatin Immunoprecipitation followed by qPCR (ChIP-qPCR) was performed with specific antibodies targeting NOTCH1 (custom made anti-serum from Covance^16^) or H3K27ac (abcam ab4729). Briefly, cells were harvested by cross linking with 1% formaldehyde for 5 (for H3K27ac) or 15 (for NOTCH1) minutes and subsequently quenched with glycine. Chromatin shearing was done in bulk with 50 – 100 million cells. Sonication was performed with a Bioruptor Pico sonicator, enriching for fragments between size 200-500 bp. Aliquots were taken so that each NOTCH1 and H3K27ac ChIP experiment used 7.5 and 2.5 million cells, respectively. Sheared chromatin was preincubated with anti-rabbit protein A agarose beads to reduce nonspecific binding. Precleared chromatin samples were then incubated with specific antibodies at 4 C overnight with gentle rotation. The next day, immunoprecipitation was performed by adding protein A agarose beads to the chromatin/antibody solution and rotating for 2 h at 4 C. Beads were pelleted and subjected to low salt (.1% SDS, 1% TritonX-100, 2mM EDTA, 20mM Tris pH 8, 150mM NaCl), high salt (.1% SDS, 1% TritonX-100, 2mM EDTA, 20mM Tris pH 8, 500mM NaCl), LiCl and TE washes, then eluted in SDS/NaHCO_3_ elution buffer. Reverse crosslinking and proteinase K (NEB) treatment were performed to release the bound DNA, which was recovered by ethanol precipitation. qPCR was performed as described in the RT-qPCR section above. Primers for qPCR are listed in Supplementary Table 1. Cq values were converted into percent input by comparing to a 1% input sample. In order to show biological replicates on the same scale, % input was normalized to WT Jurkat cell = 1 within each replicate.

### Sequence analysis

Protein sequence conservation was determined using the ConSurf server with default settings^25–28^. Predicted disorder was determined using DISOPRED3 on the Psipred server^29,30^. Amino acid charge scale was determined by assigning colors to the pI values of the amino acids.

## Author Contributions

BG, JCA, KA and SCB designed experiments, JMR, BG, and EE performed experiments, JMR, BG, and SCB drafted the manuscript, and all authors edited the manuscript.

## Acknowledgments

JMR and BG were funded by LLS CDP postdoctoral fellowships, and JMR was also supported by NIH training grant 2T32GM007748-40. This work was supported by NCI R35CA220340 to SCB. We thank members of the Blacklow and Adelman labs for helpful discussion.

---

**Supplemental Figure 1.**
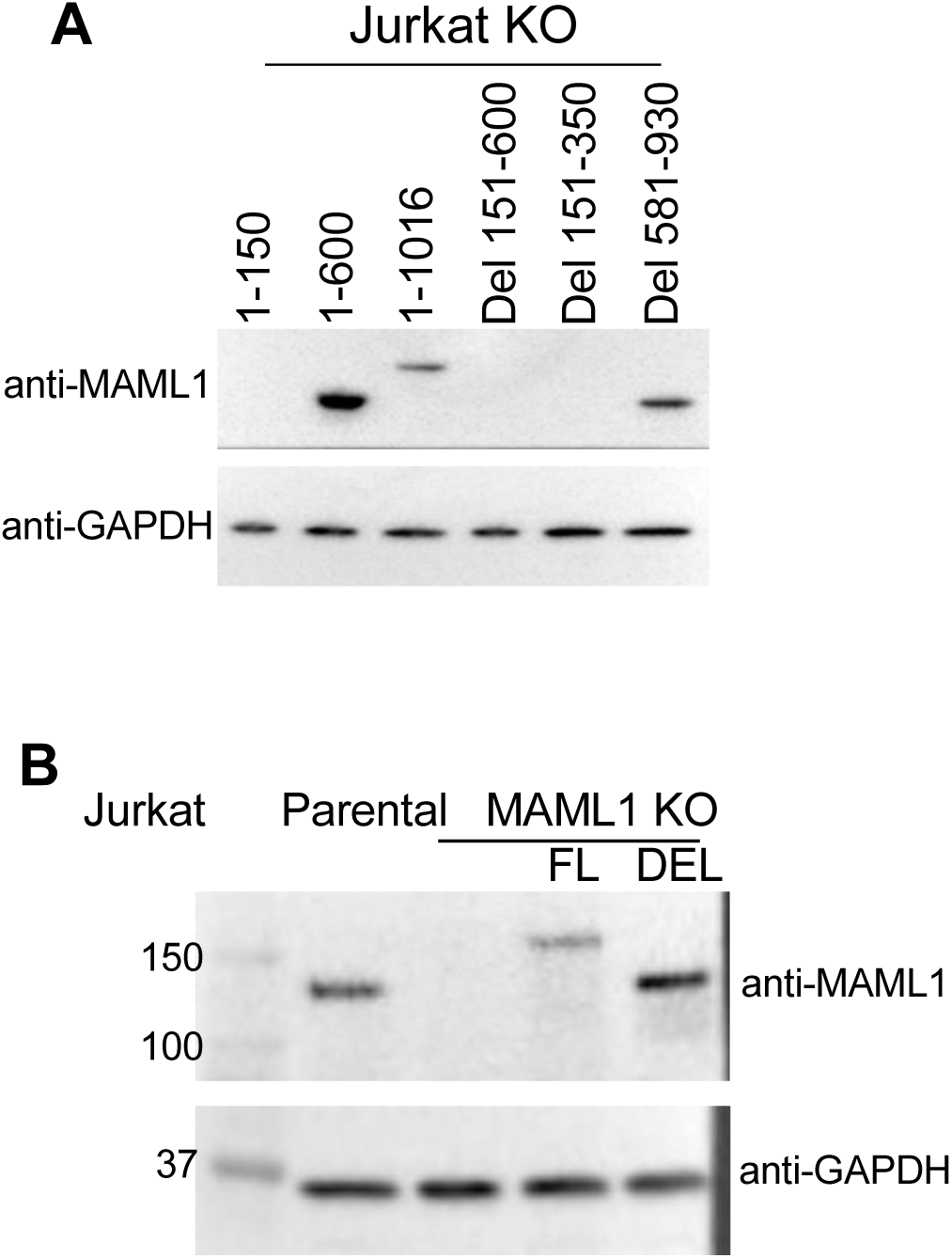
(a) Western blot of MAML1, using a MAML1 antibody that targets the N-terminus, in all add back cell lines. (b) Western blot of MAML1 in WT, KO, and FL and MAML1_Δ151-350_ add-back lines, using an MAML1 antibody that targets the C-terminus and thus will detect both FL and MAML1_Δ151-350_.

**Supplemental Figure 2.**
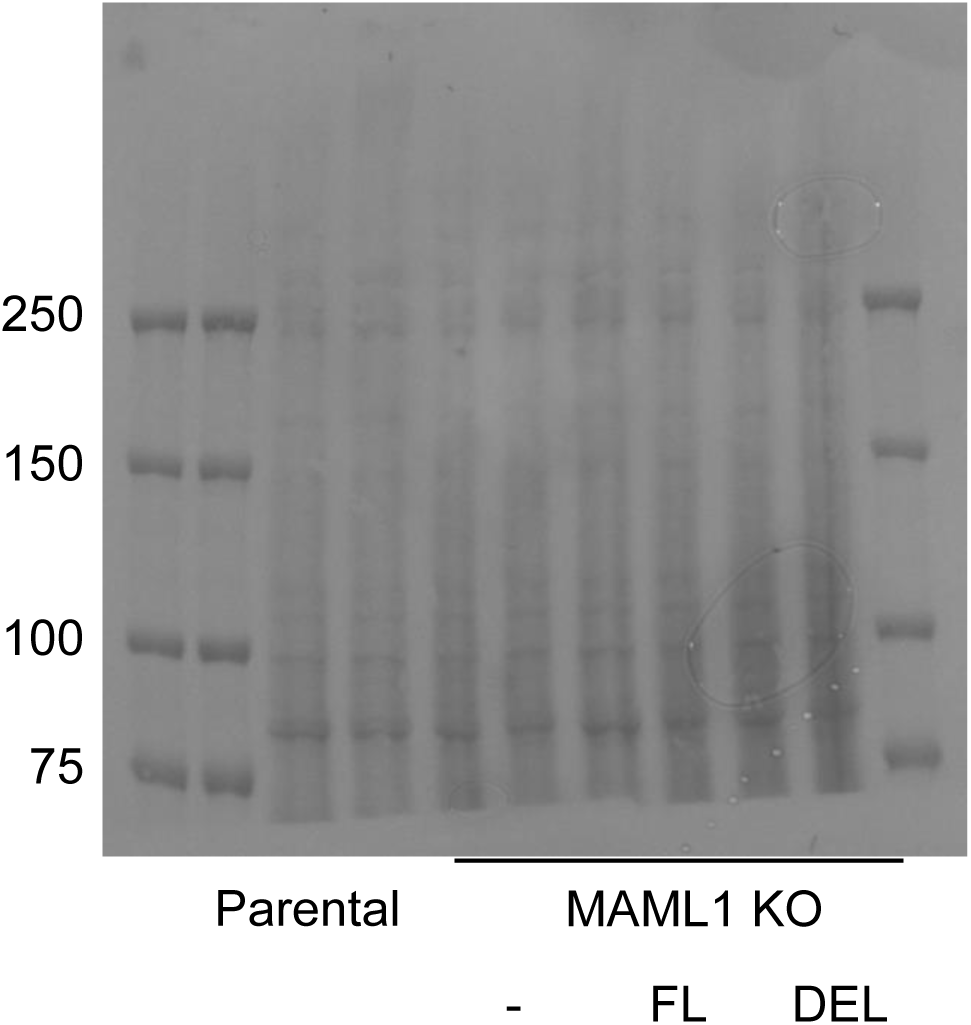
Ponceau stain of the Western Blot shown in Figure 4, used to confirm comparable protein loading.

**Supplemental Table 1**

Table of reagents used in this study.

